# Integrating single-cell and single-nucleus datasets improves bulk RNA-seq deconvolution

**DOI:** 10.1101/2025.08.20.671333

**Authors:** Adriana Ivich, Casey S. Greene

## Abstract

Bulk RNA-seq deconvolution typically uses single-cell RNA-sequencing (scRNA-seq) references, but some cell types are only detectable through single-nucleus RNA sequencing (snRNA-seq). Because snRNA-seq captures nuclear, but not cytoplasmic, transcripts, direct use as a reference could reduce deconvolution accuracy. Here, we systematically benchmark strategies to integrate both modalities, focusing on transformations and gene-filtering approaches that harmonize snRNA-seq with scRNA-seq references. Across four diverse tissues, we evaluated principal component–based shifts, conditional and non-conditional variational autoencoders (scVI), and the removal of cross-modality differentially expressed genes (DEGs). While all methods improved performance relative to untransformed snRNA-seq, filtering consistent cross-modality DEGs delivered the greatest gains, often matching or surpassing scRNA-only references. Conditional scVI performed comparably and was especially effective when matched scRNA–snRNA cell types were unavailable. In real adipose bulk samples without ground truth, DEG pruning and conditional scVI provided the most robust cell-fraction estimates across donors and transformations. Together, these results demonstrate that scRNA-seq should be prioritized as the reference when available, with snRNA-seq appended only after filtering cross-modality DEGs. For less-characterized systems where DEG information is limited, conditional scVI offers a practical alternative. Our findings provide clear guidelines for modality-aware integration, enabling near-scRNA-seq accuracy in bulk deconvolution workflows.

## Introduction

Gene expression is typically studied using bulk RNA-sequencing (bulk or bulk-RNA-seq) of a variety of tissues. Bulk RNA-seq is a relatively cost-effective way to understand the transcriptional landscape of samples in the context of cancer, cell development, infectious disease, drug response prediction, etc [1]. Bulk-RNA-seq can also be readily performed on frozen and FFPE samples, expanding the accessibility and scope of bulk data [1, 2]. Because bulk is widely available for many tissues, it can also be used to train deep learning models that require vast amounts of training data [3–5].

However, bulk-RNA-seq provides only indirect information about the cellular composition of the samples, limiting our understanding of the cell-type-specific contributions to the observed transcriptome [6]. The cell type proportions in a sample can reveal key details that influence the development of therapeutics, our understanding of the tumor microenvironment (TME), and more [7, 8]. Single-cell RNA sequencing (scRNA-seq) provides gene expression measurements of individual cells. Each individual whole cell is isolated or barcoded, and the transcriptome is sequenced [6]. ScRNA-seq gene expression counts contain both cytoplasmic and nuclear RNA, and in an ideal world, produce data directly comparable to bulk RNA sequencing.

Bulk deconvolution enables researchers to estimate cell-type proportions and cell-type-specific expression from vast archival bulk samples [9–11]. Bulk deconvolution describes the process of estimating the proportions of discrete cell types in a bulk transcriptomic sample. Accurately deconvolving bulk data could reveal changes in the TME, transcriptomic heterogeneity in response to treatments, and other key factors that could influence survival.

Reference-based deconvolution methods generally assume a strong concordance between the gene expression observed in the cell references used and the bulk at hand, so researchers often use a scRNA-seq dataset as a reference for cell type expression [12]. However, this assumption is compromised in cases where there are expression changes and notable cell losses due to the dissociation procedures or microfluidic devices used in many scRNA-seq protocols.

Apart from dissociation-specific marker upregulation [12], scRNA-seq protocols can cause cell types to be lost [13]. When cell types present in bulk are missing from single-cell observations, they substantially reduce deconvolution performance [13], making cell proportion estimates unreliable. In these cases, researchers often use single-nucleus RNA-sequencing (snRNA-seq) as a reference [14, 15] or combine both modalities [16, 17]. Cells that are sensitive or difficult to dissociate are still often observable with snRNA-seq [18–20], and cell types not often found with scRNA-seq can be seen in snRNA-seq data [20]. However, snRNA-seq captures nuclear RNA (Figure 1a), and the loss of cytoplasmic RNA compromises the assumption of high concordance.

**Figure 1.**
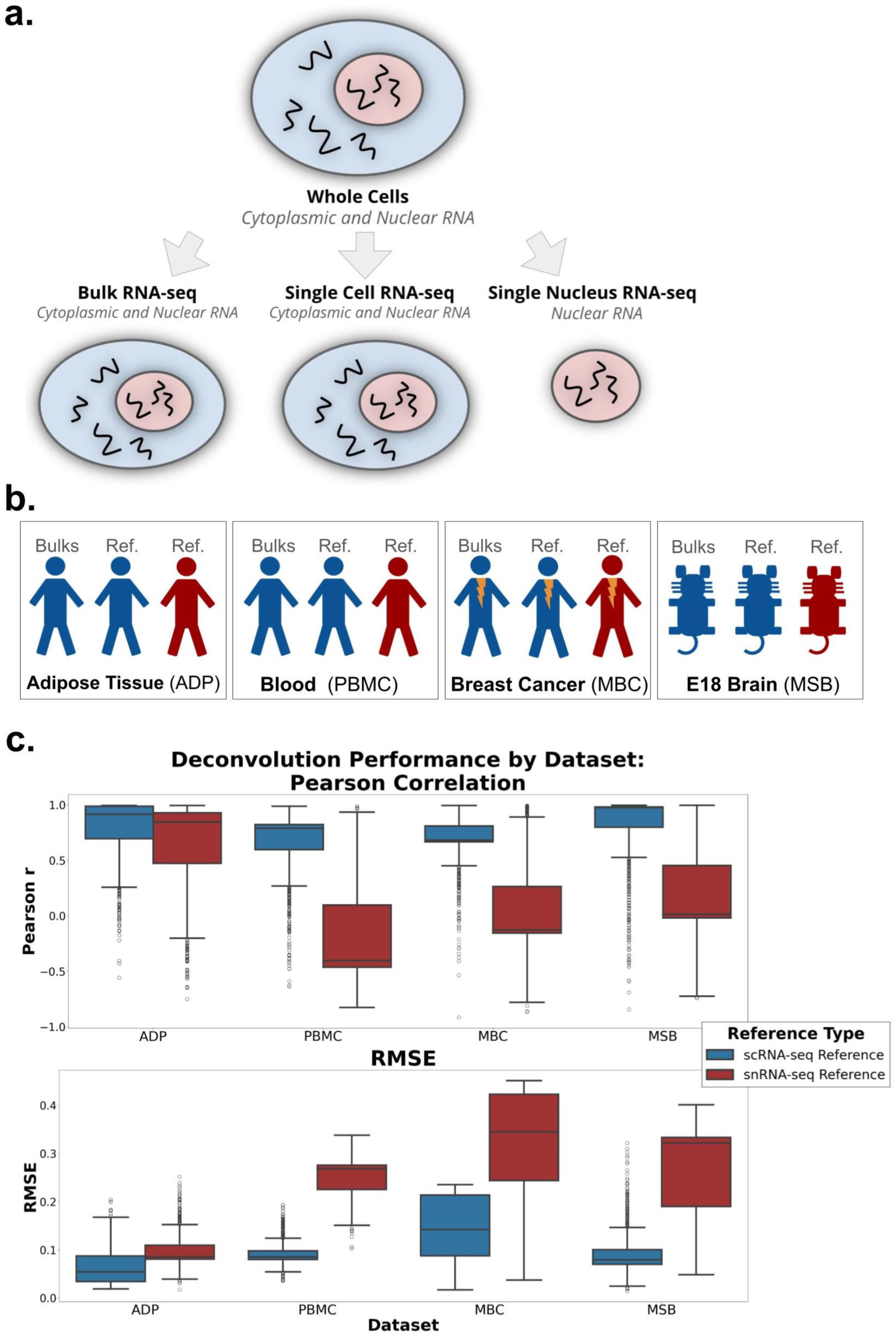
Schematics of experimental motivation and datasets used. **a** RNA sequencing yields different datasets depending on cell proportion that is sequenced. Bulk RNA-seq and ScRNA-seq) contain counts of both cytoplasmic and nuclear RNA. However, single nucleus RNA-seq contains only nuclear RNA potion. **b** Our study uses four RNA-seq dataset types: adipose tissue (ADP), peripheral blood mononuclear cells (PBMC), metastatic breast cancer (MBC), and developmental mouse brain (MSB). These datasets encompass 4 distinct and varied situations of cell loss across species. **c.** Comparison of deconvolution performance (Pearson and RMSE) of pseudobulks made from ScRNA-seq using a ScRNA-seq reference and a S-Nuc reference with same cell types. RMSE: root mean squared error. ScRNA-seq: single-cell RNA-seq. S-Nuc: single-nucleus RNA-seq.

Previous work has focused on deconvolution methods that expect or can use snRNA-seq references exclusively within certain difficult-to-assay tissues [21–23]. Other work has focused on aligning the bulk expression with the reference expression, which mitigates the gap between cytoplasmic and nuclear RNA present in snRNA-seq datasets, such as the deconvolution methods *SQUID* [24] and *BISQUE* [23]. These methods use snRNA-seq references but rely either on matched tissue (bulk and reference from the same sample), or cell proportions in the reference to match the bulk to fit the bulk expression using observed cell proportions in the reference. Both are unfeasible when the cell types references available are from different datasets or require distinct modalities to include all cell types, i.e., with unmatched snRNA-seq samples and scRNA-seq data that do not contain all cell types. Also, the constraint that bulk cell type proportions and single-nucleus proportions must match is challenging to confirm, as many tissue types can be heterogeneous in composition. While the distinction between snRNA-seq and scRNA-seq references is recognized and clearly influences deconvolution, the impact of simple transformations applied to mixed references on deconvolution performance using methods that rely on scRNA-seq is underexamined. The present study aims to investigate whether this mixed-modality reference can improve deconvolution performance in a benchmarked method that expects scRNA-seq as a reference.

In this study, we compare deconvolution performance in the context of simple transformations applied to scRNA-seq vs. snRNA-seq references. We assess gene filtering, linear principal component analysis (PCA), and non-linear scVI-based transformations [25, 26]. We benchmarked transformations on four datasets: human adipose tissue, blood, metastatic breast-cancer liver tissue, and embryonic mouse brain. We systematically substituted one scRNA-derived cell type with its (untransformed or transformed) snRNA-seq equivalent and evaluated concordance against pseudobulks, real bulk samples, and robustness across patients. While using snRNA-seq references alone substantially reduced deconvolution accuracy, simple transformations often closed the gap. In addition, filtering genes that were consistently differentially expressed between scRNA-seq and snRNA-seq cell types across tissues often improved performance in unrelated datasets to a level consistent with other transformations.

## Results

### Pseudobulk deconvolution with scRNA-seq and snRNA-seq established baseline performance

Our goal was to establish an experimental framework to compare scRNA-seq and snRNA-seq references under the assumption that snRNA-seq would represent a complete set of cell types with incomplete RNA (only nuclear), while scRNA-seq would represent the complete expression measurements for cells but for an incomplete set of types. To simulate this setting, we created pseudobulks with known proportions from an scRNA-seq sample. We then deconvolved all pseudobulks with either an scRNA-seq or snRNA-seq reference derived from a different sample of the same tissue type containing the same cell types in equal numbers. We assessed performance using root mean squared error (RMSE) and Pearson correlation of cell type proportions (see Methods for more details). We repeated this in four datasets: three human (two of reference tissue and one cancerous) and one mouse (developmental) (Figure 1b). Consistent with expectations, scRNA-seq yielded a higher Pearson correlation and a lower RMSE than the snRNA-seq reference across all datasets (Figure 1c).

We assessed the significance of differences using two-sample t-tests of Pearson correlation and RMSE across both reference types within each dataset. In the adipose dataset, the scRNA-seq reference produced significantly higher Pearson correlations than the snRNA-seq reference (t = 10.17, p < 1e-6), while RMSE values were significantly lower for scRNA-seq (t = -23.41, p < 1e-6), indicating more accurate deconvolution. Highly significant differences were observed across all datasets and all comparisons (p < 1e-6). Taken together, we found that the experimental design distinguished between positive and negative controls across all datasets.

### Multiple approaches can integrate snRNA-seq and scRNA-seq references

We implemented a set of controls and transformations (Table 1) across a set of potential algorithmic selections. These were the source modality for each cell’s expression, whether the operation was cell-type-specific or global, the transformation applied (e.g. scVI conditional, latent-shift, PCA neighbour averaging, etc.), and the final gene set used in the references.

**Table 1.**
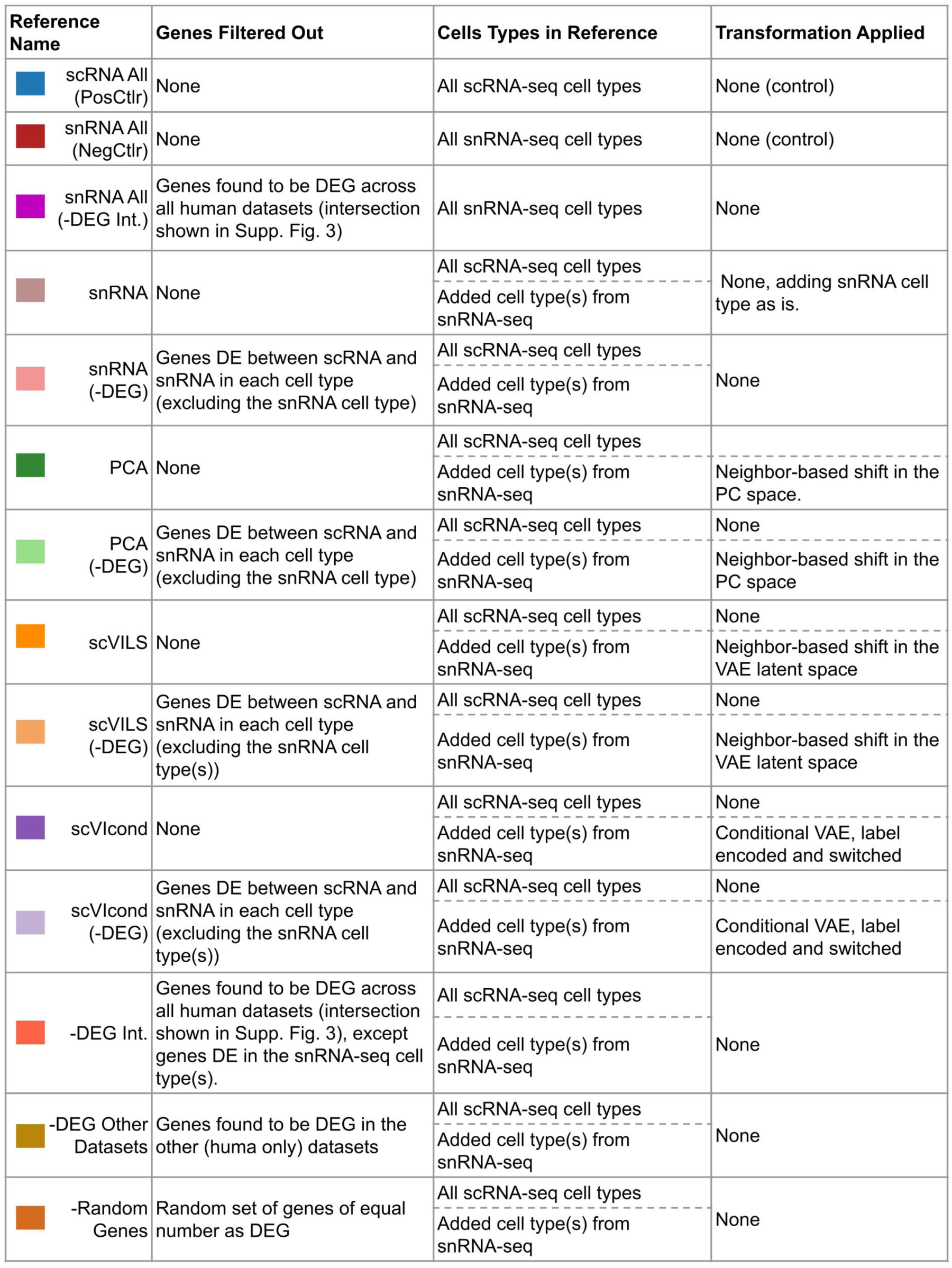
Description of each control and transformation evaluated. Reference name shows the colors used throughout the plots for each control and transformation. The table shows the reference name as is referred in the main text and figures, the origin of the cell’s expression included in the reference, whether the transformation is applied to all cells or only one cell type (snRNA-seq), the transformation type, and the genes that are present in the final reference. “All” genes included in reference (right column) refers to all genes in common between the references and pseudobulks/bulks. Note that some transformations only contain one snRNA-seq cell type, and the rest of the cells’ expression comes from scRNA-seq. The first two references listed (scRNA All (PosCtrl), snRNA All (NegCtrl)) contain all cells from only one reference with no transformation applied as controls. The snRNA All (-DEG Int.) is the only other reference type that contains all cell types from one modality, snRNA-seq. ScRNA-seq: single-cell RNA-seq. S-Nuc: single-nucleus RNA-seq.

To first test whether our transformations make snRNA-seq profiles closer to their scRNA-seq and bulk (nuclear and cytoplasmic RNA as well) counterparts, we applied the three cell transformation strategies (scVI conditional, scVI latent-shift, and PCA neighbour shifts) plus DEG-filtered variants to four datasets (details in Methods). At the single-cell level, assessing with cosine-similarity across the four datasets revealed that transforming each snRNA-seq cell type with the scVIcond model most closely aligned it to the matching scRNA-seq signature, with PCA a close second. In this context, the latent-shift adjustment offered little or no gain (Supplementary Figure 1a-d). When we aggregated the transformed fat-cell and neutrophil profiles into 100 synthetic pseudobulks and compared them with real bulk RNA-seq, PCA-based pseudobulks best resembled bulk expression, while scVIcond-transformed pseudobulks were least similar; after removing cell-type-specific DEGs, the highest bulk-level similarity was achieved simply by using DEG-filtered snRNA-seq, followed by the PCA-DEG transform (Figure S1e-f). Together, these results indicated that scVIcond excelled at aligning individual nuclear profiles to single-cell references, whereas PCA (especially with targeted gene filtering) produced the most bulk-like composite expression.

### Transforming snRNA-seq measurements in an scRNA-seq references improves deconvolution

We then evaluated the performance of deconvolution with ground truth proportions. For each tissue type (Figure 1b), we used 3 samples: one scRNA-seq to create pseudobulks, one scRNA-seq for reference, and one snRNA-seq used to apply and test our transformations. We created pseudobulks with the scRNA-seq designated sample with all cell types with more than 50 cells (see Methods for details of pseudobulk creation). We created two control references for deconvolution for each tissue type: one with all cells from the reference scRNA-seq sample (scRNA All (PosCtrl)) and a second with all cells from the snRNA-seq sample (snRNA All (NegCtrl)).

In each, we held out one cell type at a time from a scRNA-seq sample and replaced that cell’s expression with an equivalent from the snRNA-seq sample. This snRNA-seq cell type was either added as is (snRNA) or with each of our transformations (PCA, scVIcond, scVILS, and the same without DEGs; see Table 1 and Methods for more detail). All the transformations contain the same cell type’s expression, just from a different data modality source. We hypothesized that the transformations would improve the deconvolution performance when compared to the snRNA All (NegCtrl), and closer in performance to the scRNA All (PosCtrl) reference. This positive control reflects the ideal but unexpected scenario where all cells available in the bulk are also present in scRNA-seq.

We evaluated the deconvolution in three different groups, each corresponding to the specific cell type proportions of interest. In the first group, “All Cells”, we evaluated the performance in estimating the cell type proportions of all cell types in the pseudobulk, independent of held out status. The second group, “Non-Removed Cells” evaluated performance only on the subset of cell types that were not removed, assessing the impact on cell types that were present in scRNA-seq data. Lastly, we subset performance to “Removed Cell Only” to examine the accuracy of proportion estimates for the cell types that were absent in the scRNA-seq data.

We grouped all performance metrics across the four tissue types (Figure 2: dot = mean; bars = 95% bootstrapped confidence interval (CI) of the mean, 1000 iterations). For “All Cells” (Figure 2a), the scRNA All (PosCtrl) achieved the highest Pearson correlation, followed closely by the snRNA -DEG transformation and then scVIcond transformation. The remaining methods performed somewhat comparably, but much better than the snRNA All (NegCtrl). Using all cells from a snRNA-seq sample as reference yielded the worst performance in both RMSE and Pearson correlation metrics, with the upper 95% CI failing to exceed a correlation higher than 0.2. Surprisingly, the snRNA-DEG transformation outperformed the positive control; in fact, the positive control’s RMSE was comparable to that of PCA -DEG. Notably, removal of DEGs boosted performance in all transformations (e.g., scVIcond -DEG achieved higher Pearson correlation and lower RMSE than scVIcond).

**Figure 2.**
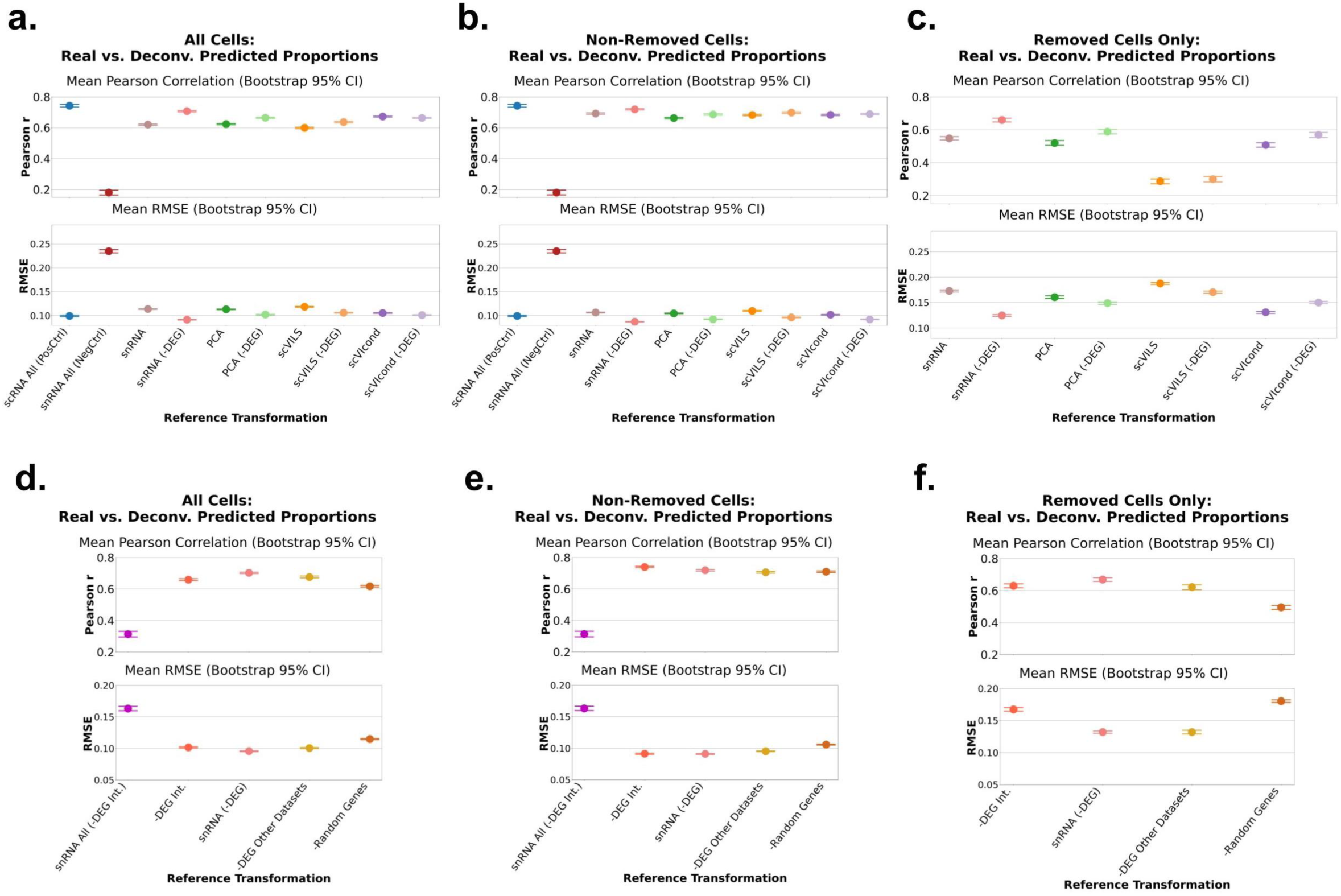
Pseudobulk deconvolution accuracy with each cell type in scRNA-seq held out and transformed. Each plot shows the Pearson correlation value (top panels) and the RMSE values (bottom panel) for the ground truth pseudobulk (simulated) proportions and the predicted proportions. We hold out one cell type at a time from each scRNA-seq dataset and replace that cell type’s expression with a snRNA-seq equivalent with each of the transformations or controls (scRNA All and snRNA All) on the x axis of each plot. Each dot represents the mean metric (correlation or RMSE) across datasets (Figure 1b) and the bars represent the 95% bootstrapped confidence interval of the mean. We evaluated 3 scenarios (see Methods for details). a and d. All cells included in performance metrics calculations, b and e. Non removed cells only included in performance metrics calculations and c and f. Only removed cells included in performance metrics calculations. Per dataset metrics can be seen in Supplemental Figure 2. Y-axes are truncated to highlight the variation. scRNA-seq: single-cell RNA-seq, RMSE: root mean squared error, snRNA-seq/snRNA: single-nucleus RNA-seq.

Across tissue types, these trends held generally. The per tissue deconvolution performance is shown in Figure S2. Notably, the negative control reference in the ADP dataset had surprisingly good performance (mean ≈ 0.65), although it dipped as low as -0.2 in other datasets. Nevertheless, it exhibited the lowest performance in all settings.

In the “Non-Removed Cells” scenario, the overall patterns persisted: the positive control again led in Pearson correlation, and the negative control correlation remained below 0.2 (Figure 2b). In RMSE, several DEG-removed methods (snRNA -DEG, PCA -DEG, scVIcond -DEG) again outperformed the positive control, and the performances of scVIcond and scVILS -DEG were comparable. Per dataset analysis mirrored the “All Cells” results, with the ADP dataset showing particularly high Pearson correlation and low RMSE across methods compared to the other datasets yet still ranking the negative control worst within that sample (Figure S2b).

The “Removed Cell Only” scenario is unevaluable with the positive and negative controls, since those references didn’t include removed cells. In this scenario we observed the largest inter-transform variability in the mean Pearson correlation and RMSE (Figure 2c), hinting at cell-type-specific advantages to each transform. This high transform-to-transform variability was also observed at the dataset level (Figure S2c). SnRNA -DEG achieved the highest Pearson correlation and lowest RMSE (Figure 2c). The PCA -DEG and scVIcond -DEG transformations yielded Pearson correlation values higher than the positive control, and all transformations (except scVILS) achieved lower (or comparable for scVILS -DEG) RMSE. At the per-dataset resolution, the scVIcond mean Pearson correlation was greatly decreased by the PBMC dataset, and without considering this dataset the scVIcond would outperform all other transform’s Pearson correlations (Figure S2c). This could suggest that the conditional variational autoencoder (VAE) model for the PBMC dataset might require different training parameters or datasets. The MSB dataset exhibited uniformly poorer RMSE across all transforms (Figure S2c).

Together, these findings revealed that removing DEGs (-DEG) was sufficient to improve deconvolution performance across many scenarios and datasets. The DEG-pruned transformations, especially the snRNA -DEG, enhanced deconvolution accuracy across all evaluation scenarios, often in line with positive control and deep learning performances.

### Selective pruning of dataset-specific and cross-dataset DEGs improves deconvolution accuracy

We then aimed to test whether removing DEGs observed between modalities across multiple datasets was sufficient to improve deconvolution performance. This would avoid the need to have matched snRNA-seq and scRNA-seq data from enough participants to have well-powered tests of differential expression. We considered that DEG between cell types observed in multiple other tissue datasets (human only) could enable the filtering of genes that translate across tissue types (-DEG Other Datasets). We also removed the intersection of genes classified as differentially expressed in all three human datasets in at least one cell type (-DEG Int.). This list of genes is described in Figure S3. Moreover, as a negative control we tested removing a random set of genes matched in size to the -DEG set to evaluate whether simply removing genes or features improved performance. Lastly, to test whether gene pruning would be sufficient to obtain good performance for a full snRNA-seq reference, we also tested a full snRNA-seq reference without the DEGs found across all human datasets (-DEG Int.). All pruned gene lists and references created are summarized in Table 1. We evaluated the impact of these gene removal strategies on deconvolution accuracy across the three evaluation scenarios (“All Cells”, “Non-Removed Cells and “Removed Cells”).

In the “All Cells” scenario, pruning the dataset specific DEGs (the same as snRNA - DEG, as above) again delivered the best performance, achieving the highest mean Pearson correlation and the lowest RMSE (Figure 2d). By contrast, the pruned full snRNA-seq reference snRNA All (-DEG Int.) produced the poorest accuracy, with the lowest correlation and highest error. Removing DEGs defined in other human datasets (-DEG Int.) yielded results that were close to the best case, indicating that simply filtering genes commonly differentially expressed between modalities across tissues recovers much of the benefit of dataset-specific DEG pruning.

When focusing in the “Non Removed Cells” scenario, -DEG Int. yielded the highest Pearson correlation and an RMSE on par with the snRNA -DEG reference (Figure 2e), suggesting this genes list integrated across tissue types robustly reflects scRNA-seq and snRNA-seq differences. The random gene control showed the highest RMSE again, and surprisingly had comparable Pearson correlation mean to the -DEG of the other dataset’s removal. Notably, the combined DEG list from other datasets was quite large, reducing the reference to only ∼6,000 genes in some cases. This may have discarded biologically informative markers needed for optimal deconvolution, and there are likely more sophisticated strategies to optimize filtering.

The results are comparable to “All Cells” otherwise. It is worth noting that in the adipose dataset only, snRNA All (-DEG Int.) performed better than -Random Genes and -DEG Other Datasets (Figure S2d-e). We attribute this higher performance in the adipose dataset to this tissue type having fewer compartmental biases, i.e., less dramatic nuclear-to-cytoplasmic transcript variation (Supp. Table 1). Of all tissues tested it shows the lowest total number (and percentage) of DE genes, hinting at fewer nuclear to cytoplasmic transcript differences.

In the “Removed Cells” scenario, the random gene removal again performed as hypothesized, showing the lowest Pearson correlation and highest RMSE (Figure 2f). Interestingly, pruning the identified DEGs from other datasets produced the highest Pearson correlation of all strategies but was outperformed in RMSE by snRNA -DEG (Figure 2c). The DEGs of a specific dataset (snRNA -DEG) only includes those of the cell types that are not removed, mimicking a real-life situation. The boost in performance in the “Removed Cells” scenario by -DEG Other Datasets suggests that removing genes with consistent differential expression across tissues may better generalize to recovering an absent cell type. Overall, these findings reinforce that gene set pruning, especially of dataset specific or cross dataset DEGs, can significantly enhance deconvolution when reference panels are incomplete, whereas indiscriminate gene removal offers little to no benefit. The performance metrics per dataset are shown in Figure S2 (d-f).

Due to the high performance of the -DEG Int. (i.e., genes that were found to be differentially expressed in at least one cell type in all three human datasets), we also analyzed the biological component involved with this list of genes. We hypothesized that the improvement in deconvolution performance by removing these genes is due to the high concordance of this list with intrinsic variations in cytoplasmic and nuclear RNA (i.e., scRNA-seq and snRNA-seq). We performed gene ontology (GO) analysis (see Methods for details) and found the statistically significant components were related to cytoplasmic or ribosomal components of the cell (Figure S3b). This again suggests this list could have biological meaning and be applicable to other datasets. In Table 2 we include a table summary of the number of DEGs per cell type across all datasets, the percentage of total genes these DEGs constitute, and the number of genes in common between cell types for each of the datasets used, including those cell types in common.

### Methodological robustness differentiates transformation strategies

We next aimed to test the transformed reference expression while deconvolving real bulks. Real bulk data does not have a reliable ground truth for proportions, so each reference yields arbitrary predicted proportions. Therefore, we used consistency in predictions as a measure of robustness, similar to previous work [27]. The idea is that if a transformed reference yields similar predicted proportions as the other transformations, it is likely to be reliable and robust transformation [27]. We employed a real-setting experimental design where we used all cells observed in 7 samples of scRNA-seq adipose tissue, and two cell types which are not observed in any scRNA-seq sample were added from 12 snRNA-seq samples. We transformed these 2 “missing” cell types, fat cells and neutrophils, with each of the transformations, and added the transformed snRNA-seq cell to the scRNA-seq observed cell types to create one reference per transform, so all scRNA-seq cells and transformed snRNA-seq. This is similar to what was done in the pseudobulks vs. real bulks experiment above (see Methods for details).

We used each of these references (i.e., with each of the transformations) to deconvolve 434 real bulk samples. We compared the predicted proportions of each used reference with the predicted proportions of the other references (one vector of all predicted proportions vs. one vector of all predicted proportions) and computed the cosine similarity per transform pair.

The transform-to-transform similarities are summarized in a heatmap in Figure 3a. Overall, most transformations had high concordance in predicted proportions with each other, except for the PCA -DEG reference. We computed the 95% CI of the mean cosine similarity per transform and show these distributions in Figure 3b. Surprisingly, removing the “intersection genes” (-DEG Int.) i.e., the set of genes found to be differentially expressed in all three human datasets, had the largest mean cosine similarity to the other transforms, reiterating the potential power of integrating the -DEGs across tissue types.

**Figure 3.**
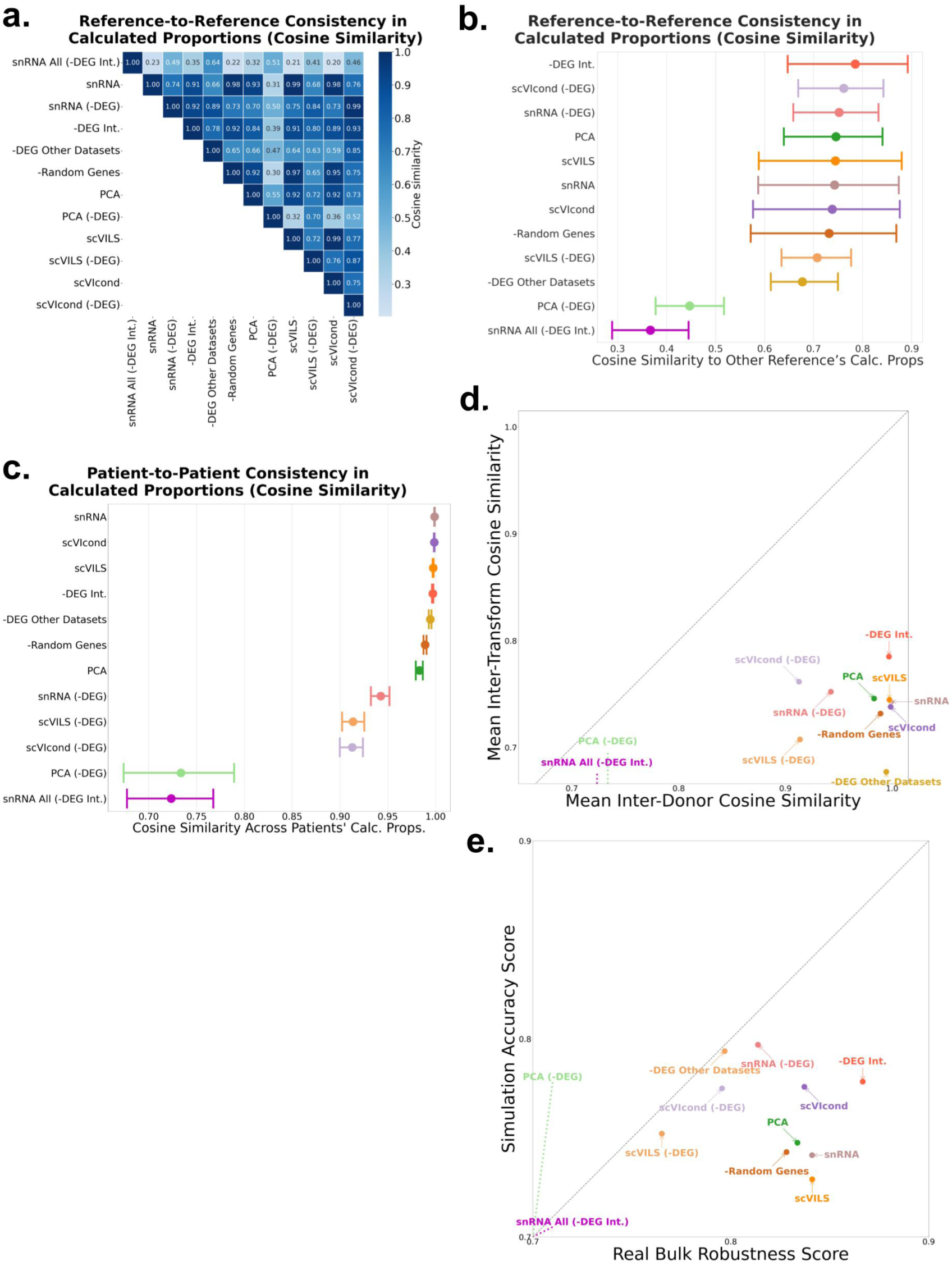
Evaluation of real adipose bulks deconvolved with each transformation and final scores per transform. **a** Heatmap showing the cosine similarity of calculated proportions per transformation - similarity between calculated proportions indicates reliability. b. The mean (dot) cosine similarity per transform, shown in a, with 95% bootstrapped confidence intervals. Transforms ordered by mean value. c. Cosine similarities between calculated proportions using fat cells from different patients per transform, the mean value as a dot and the 95% bootstrapped confidence intervals shown. Transforms ordered by mean value. d. Composite robustness and accuracy scores including inter-transforms and inter-patient similarities and RMSE and Pearson correlation values respectively. The transforms in the upper right quadrant have high accuracy and robustness indicating reliability. X and y-axes are truncated to highlight the variation. The transforms that have the highest value across both axes are considered the yield the most robust and accurate results. scRNA-seq: single-cell RNA-sequencing, RMSE: root mean squared error, snRNA-seq: single-nucleus RNA-sequencing, DEG: differentially expressed gene.

We then explore another form of robustness by using cells from a different patient in each reference and comparing similarity in predictions. Similarly, we expect that high concordance in the predicted proportions between different patients’ cell types would signify high robustness in the transformation. We used snRNA-seq-derived adipocytes from 12 patients individually (same data as above, just not combined) and transformed these cells with each of the described transformations. We used this reference per patient, per transform, to deconvolve the same 434 bulk samples (same data as above). We calculated the cosine similarity, as above, for each transformation across the predicted proportions of each patient’s reference.

Most transforms achieved comparable cosine similarity (scVIcond, -DEG Int., scVILS, - DEG Other Datasets), with snRNA all (-DEG Int.) and PCA -DEG achieving similar worse similarity (Figure 3c).

Interestingly, adding raw snRNA-seq adipocytes to the reference yielded the highest mean (although comparable to the other transforms) of cosine similarity across all transformations. Also, all the transformations that include the removal of DEG had the lowest patient-to-patient concordance (Figure 3c). We observed in the snRNA-seq data that not all cell types appear in every donor, so each donor’s DEG list is based on a different subset of cell populations. As a result, the genes we remove can differ wildly in identity and number across donors making each donor specific reference diverge rather than align.

We then combined both robustness measures; inter-transform (proportions predicted between transforms using same patient data) and inter-patient (proportions predicted between patients using same transform). We plotted each in an x and y axis respectively, expecting the most robust transformation to have high values across both axes. We found one cluster of the most robust transforms that includes, in order: -DEG Int., scVILS, scVIcond, snRNA-seq and PCA. Surprisingly, the removal of random genes (-Random Genes) had high values across both axes too. We discuss this below.

### Joint accuracy-robustness scoring highlights well-performing transformation strategies

We then evaluated each of the transformations in a way that combined both the deconvolution performance observed in the simulation studies (i.e., pseudobulks with ground truth) and the robustness observed in the inter-transform and inter-patient consistency. To enable comparison, we integrated results for each transform into two scores; one that summarizes the RMSE and Pearson correlation values observed in all simulation studies (i.e., all cell types in all datasets), and one that summarizes the consistency in calculated proportions (real bulks) from the inter-transform study and the inter-patient study.

We normalized the RMSE and Pearson correlation values within each dataset by min-max scaling them to the range [0,1] (after inverting RMSE so that higher values indicate better performance). These normalized scores were then averaged across all samples for each transform, and subsequently averaged across datasets to obtain a single normalized RMSE and normalized Pearson correlation per transform. We then took the mean of these two values to obtain the composite “simulation accuracy score.” Similarly, we normalized and averaged both the cosine similarities per transform and per patient, yielding a composite “robustness score” for each transform (see Methods for details) (Figure 3e).

We observed the removal of the intersection genes, -DEG Int. have the highest robustness score, and the third highest accuracy score overall, further suggesting this list of genes reflects true snRNA-seq to scRNA-seq differences that can aid deconvolution if removed from the reference. The scVIcond also has a high score across both axes, making it a very competitive alternative. The removal of the DEGs from the dataset (snRNA -DEG) had the highest accuracy, but because of the high donor to donor variability, it has the second worst robustness score, second to only the -DEG Other Datasets reference. We hypothesize again that this is due to the high data to data variability, and quantifying a different set of DEGs per dataset could give vastly different results depending on the number of matched cells, etc.

The snRNA All (-DEG Int.) reference had the worst score in both accuracy and robustness. This highlights that even if the pruning of the -DEG Int. is shown to be improve performance in mixed references (snRNA-seq added to scRNA-seq), it is not a viable alternative when using a full snRNA-seq reference.

Surprisingly, the removal of a random set of genes performed better than some of the more involved transformations. This likely reflects the fact that most of a cell’s identity is encoded in a relatively small core of highly informative marker genes, or “cell markers”. By randomly dropping a fraction of features we might almost never eliminate those key genes. Instead, it is more likely we filter out a large amount of low-expressed, noisy transcripts. That uniform “feature dropout” could act like a lightweight feature filtering step, helping to stabilize both inter-patient and inter-transform comparisons. An alternative explanation is the high collinearity observed in scRNA-seq data, so many genes can act as interchangeable markers of the same underlying regulatory program. Thus, random feature dropout could remove mostly redundant information.

## Discussion

The typical workflow for a researcher aiming to infer cell type proportions of bulk samples is to get a scRNA-seq sample (either new or available online) to use as reference. If all cell types expected in bulk samples are observed in scRNA-seq, this workflow is expected to yield reliable proportions, yet the field now recognizes that this assumption often fails [13, 24]. *Maden et al.* showed that scRNA-seq and snRNA-seq capture different slices of the transcriptome, and that mismatch alone can skew cell-fraction estimates [28]. They recommended doing controlled tests where the same tissue samples are profiled with both scRNA-seq and snRNA-seq, then deconvolved side-by-side, so that any errors caused by the protocol difference become obvious [28]. In the case where some cell types are missing, which we hypothesize is often, researchers have avoided this inconsistency by either using only snRNA-seq, or scRNA-seq combined with snRNA-seq, to fill in the gap [12, 16, 19, 24]. Another workaround has been building tissue specific deconvolution models, such as *scNucConv* [21] and *DeTREM* [22] for brain and human subcutaneous adipose tissue, respectively. However, these do not generalize beyond the tissues they were built for: even visceral adipose tissue had decreased performance for *snRNAConv* built for subcutaneous adipose tissue [21].

*SQUID* [24] and *BISQUE* [23] can use either snRNA-seq or scRNA-seq as reference and attacked the same problem from the bulk side: they leave the single-cell reference as is and instead mathematically reshape each bulk RNA-seq profile based on the reference. Our study breaks key assumptions the methods rely on, namely that the bulk and single cell profiles be derived from the same patients or have the same cell type distribution, so we were unable to fairly evaluate them. Our study is the first to study the integration of both modalities from different participants in a single reference, and tackles the complementary reference side of the bias coin: we show that snRNA-only references are inadequate across four diverse tissues, but that blending snRNA-seq with scRNA-seq restores and sometimes surpasses scRNA-only performance once you prune protocol-specific DEGs or apply deep learning-based transforms (e.g., scVIcond). Together, these works outline a full toolkit: adjust the bulk when the reference is trustworthy (*SQUID/BISQUE*); adjust or augment the reference when it is incomplete or protocol-mixed (our approaches, which can be used with any deconvolution method) and fall back on tissue-specific nuclei models only when neither option is feasible.

Our findings offer a clear cautionary tale: snRNA-seq references are not interchangeable with scRNA-seq references in deconvolution workflows. The substantial decrease in performance observed by using snRNA-seq as reference vs. scRNA-seq, even with the same cell types in the same number, is a clear warning to researchers to refrain from using snRNA-seq as deconvolution references. We advise researchers to exclusively use a scRNA-seq reference for the cells that are available in scRNA-seq datasets, and when needed, append snRNA-seq cell types to the reference only using one of the transformations described in this study and not “as is”. Even in the cases where the snRNA-seq cells are not of interest to enhance deconvolution accuracy overall, we recommend including snRNA-seq-exclusive cell types in the reference.

We did not find a transformation that outperformed all others in all datasets, across all accuracy and robustness metrics, and the recommended transformation will depend on cell types present (scRNA-seq vs. snRNA-seq) and computing power available. We believe the heterogeneity between tissue types, cell types observed, tissue processing and cell number will make a “one size fits all” transformation a complicated endeavor. One key consideration in the choice of transformation is the data available: some transformations don’t rely on matching cell types across scRNA-seq and snRNA-seq datasets (PCA, scVIcond, scVILS), making them a possible solution in the cases where scRNA-seq and snRNA-seq observed cell types do not match. When the tissues being sequenced are small and there is only one sample for each modality, it could be difficult to find more than one matching cell type. The transformations that depend on matched cell types, namely the calculation and removal of dataset specific DEGs, could be a viable solution when researcher have multiple batches and thus increased observed cell number. We also separated the deconvolution accuracy metrics by scenario (i.e., All Cells, Non-Removed Cells, and Removed Cells) (Figure 2) and by dataset (Figure S2) to encourage researchers to find the transformation with the best performance in the most relevant tissue type and scenario.

We found that including raw snRNA-seq adipocytes in the reference achieved the highest average patient-to-patient cosine similarity across transformations, though the improvement was small compared with other transforms. We hypothesize that this is because nuclear RNA tends to be enriched for unspliced, nascent transcripts or “housekeeping” gene programs [29], which typically have low cell-to-cell or donor-to-donor variability, making a donor-to-donor comparison unexpectedly favor an unchanged snRNA-seq transcript. In contrast, our neighbour-based and VAE based transforms likely accentuate biological and technical differences between batches, and thus, between donors, so despite potentially improving deconvolution accuracy, this lowers the apparent patient-to-patient consistency.

A computationally reasonable alternative is removing the DEGs between scRNA-seq and snRNA-seq cell types, which made the biggest distinction in deconvolution performance across transformations in the simulations. In fact, adding snRNA-seq cell to a scRNA-seq reference and just removing DEGs with no other transformation outperformed some more computationally expensive transformations in some cases. However, this is not a one-size fits all; we observe low donor to donor consistency in all transformations involving the removal of DEGs in real bulks.

The identified DEGs greatly depend on which cell types are observed in the scRNA-seq and snRNA-seq datasets, since we calculate DEGs per matched cell type. This makes the removal of DEGs a less dependable strategy and should only be used in the cases where there are at least 4 cell types in common between snRNA-seq and scRNA-seq, and these cells are observed in reasonable numbers of more than 50. These are the parameters used in our simulations, showing high efficacy. Further work is required to evaluate whether fewer cell types or cell examples is a viable alternative.

If enough computing power is available, we recommend using scVIcond (i.e., a conditional implementation of the scVI VAE) as the first strategy. This transformation had high accuracy in simulations and high robustness (Figure 3e). Additionally, it does not depend on scRNA-seq and snRNA-seq having cell types in common for training, making it a feasible option in small datasets.

Our results indicate that removing the intersection of DEGs between 3 tissue types offers the most cost effective path to the greatest deconvolution accuracy gain, making it the preferred strategy for integrating scRNA-seq with snRNA-seq samples. We found that -DEG Int. scored the highest consistency score, and the second highest accuracy (second to only -snRNA -DEG, which is third last in consistency). These high scores, along with the gene ontology components observed, suggests that integrating multiple tissue types in the DEG analysis can further enhance the effect of scRNA-seq vs. snRNA-seq DEGs removal. We have included this list of genes in the GitHub repository for download and use [30]. We encourage researchers to use and add onto this list with their own tissues of interest, which we hypothesize will further enhance the efficacy of this list. Nevertheless, this list is only applicable to human datasets, and further work is needed to find and validate an equivalent in other species.

### Limitations of the study

Several limitations point the way forward for future studies. First, the removal of DEGs between scRNA-seq and snRNA-seq cell types, although powerful and simple, could be enhanced further by including more datasets and cell types and optimizing thresholds. We also did not protect cell type marker genes, which might provide finer control. Second, we used scVI transformation types (scVILS and scVIcond) with hyperparameters previously observed to perform well on alignment tasks [25], but further tuning per tissue type could further enhance their accuracy. Third, the space of possible transformations (e.g., non-linear domain-adaptation nets, contrastive embeddings, cross-modal diffusion models) is still largely unexplored, and methods purpose-built and hyper-parameter-tuned for this task may yet outperform the transformations we explored.

These results underscore the non-interchangeability of these modalities in deconvolution but also pave the way for future studies to compare the two in other scRNA-seq-targeted methods other than deconvolution (e.g., cell type annotation, batch normalization, deep learning models, etc.) rather than assuming snRNA-seq will perform equivalently.

Additionally, we tested only four tissues, and although the tested tissue types encompass a wide range, additional modality mismatches could be observed in other tissues that are hard to anticipate. Finally, our study focuses only on one deconvolution method, and while we do not expect results to vary greatly with other standard methods, future work should explore data integration in different deconvolution methodologies. We have included instructions on how to include a new dataset or transformation method in the GitHub repository [30] to allow researchers to quickly perform their own analyses. Together, we aim for our open-source framework and modality-aware guidelines to give the community a practical roadmap for turning vast bulk RNA-seq archives into cell-resolved insights.

## Author contributions

Conceptualization, A.I and C.S.G.; methodology, A.I and C.S.G.; software, A.I and C.S.G.; formal analysis, A.I and C.S.G.; data curation, A.I and C.S.G.; visualization, A.I and C.S.G.; resources, A.I and C.S.G.; writing - original draft, A.I and C.S.G.; writing - review & editing, A.I and C.S.G.; funding acquisition, C.S.G.

## Declaration of generative AI and AI-assisted technologies in the writing process

During the preparation of this work, the authors used ChatGPT (GPT-3-turbo) to improve the grammar and flow of the paragraphs. After using these tools, the authors reviewed and edited the content as needed and take full responsibility for the content of the publication.

## Methods details

### Datasets

This study utilized exclusively publicly available datasets. The datasets used for the pseudobulk simulation experiment consist of one snRNA-seq and two scRNA-seq batches (see Data Availability section for access details). To ensure a diverse and representative sample, we used a variety of tissue types with variable dissociation-induced bias or cell loss from diverse sources. Specifically, we used data from two non-pathological human tissues: one with dissociation bias based on prior evidence (adipose tissue) and one with minimal hypothesized dissociation bias (PBMC tissue) from 10x Genomics. We also used a cancerous tissue, metastatic breast cancer, which have significant differences compared to non-pathological tissues. Lastly, we included a E16 (in development) mouse brain tissue to see the applicability in datasets from nonhuman species and in early-stage differentiation.

Each of the datasets consists of 3 batches: one scRNA-seq dataset to create pseudobulks, an independent (not same patient) scRNA-seq dataset for single-cell references, and one snRNA-seq reference to add to scRNA-seq deconvolution reference. In the case of the MBC, the 2 reference datasets come from the same patient. For the ADP, PBMC and MSB data, the two reference datasets come from different patients/mice and protocols. In all cases, the reference datasets and the pseudobulk reference are not the same patient.

For the real bulk analysis, we used 434 real bulk RNA-seq samples, and an additional 12 patient samples of snRNA-seq and 7 samples of scRNA-seq.

### Cell type assignment

For both the ADP and MBC datasets, cell types were assigned in the original publication, and those cell types were used for the analysis. For the PBMC and MSB datasets, we assigned cell types using CellTypist [31], a logistic regression based assignment to ensure consistency and reproducibility. We used the “developing mouse brain” v1 [32] model for MSB and the “Healthy COVID19 PBMC” v1 models [33] for PBMC.

### Data preprocessing and filtering for pseudobulks and references

All datasets used are publicly available and previously processed. We added a file to the GitHub repository [30] that contains details and links for all data sources for each dataset used in this study, as well as any additional filtering parameters when needed.

For each data type (ADP, PBMC, MBC and MSB) we removed all cell types with less than 50 cells in either of the datasets to ensure sufficient variability for pseudobulks and references in the simulation studies. We also aligned the gene expression matrices of the scRNA-seq and snRNA-seq datasets to include only the common genes between them.

### Deconvolution of pseudobulks using references with transformed held-out cell types

After data and cell type filtering, we created pseudobulks with known proportions and deconvolved them with various reference types. This process was the same for all datasets (ADP, PBMC, MBC and MSB). We created pseudobulks with all available cell types using one of the scRNA-seq datasets. In order to mimic what happens in a true research setting, we removed one cell type at a time (“held-out”) from each of the scRNA-seq references and replaced this cell type with the same cell type but from snRNA-seq (nuclear RNA only) either “raw” (no change to the data) or with different transformations to harmonize the expression profiles of the held out snRNA-seq cell type with the rest of the scRNA-seq reference (see below). We created two control references: one with all scRNA-seq cells (positive control), and one of all snRNA-seq cells (negative control), both matching all cell types present in the pseudobulks. For all references and in all transformations, the number of cell examples per cell type was kept the same.

For each held-out cell type, we trained a separate scVI model that excluded that cell type from both the scRNA-seq training set and the PCA fitting step. This approach mimics real-world scenarios in which certain cell types are missing from the reference dataset, ensuring that all transformations remain valid and that our results closely reflect practical performance.

### Pseudobulks

We created pseudobulks for each of our datasets (ADP, PBMC, MSB, and MBC). All pseudobulks only contain scRNA-seq cells from the pseudobulk-dataset, not used in the scRNA-seq reference and independent of the scRNA-seq reference (i.e., not same patient). We used a custom sampling strategy to create pseudobulks of known proportions. For each pseudobulk sample, a cell-type proportion vector was generated according to one of two schemes:

- **Random Proportions:** A Dirichlet distribution (with equal concentration parameters) was used to generate a random proportion vector. In cases where the resulting cell counts (i.e., the product of the proportion vector and a fixed number of cells 1,000) resulted in zero for any cell type, the sampling was repeated until every cell type was represented.
- **Realistic Proportions:** The empirical observed cell-type proportions (as computed from the scRNA-seq data) was used as the base proportion, with a small normally distributed noise (mean = 0, SD = 0.01) added. The resulting noisy vector was normalized to add up to 1, and cell counts were computed as described for the random case. Again, resampling was performed if any cell type was assigned zero cells.

For each pseudobulk sample, the required number of cells from each cell type (as determined by the proportion vector) was randomly sampled from the scRNA-seq data without replacement when possible (i.e., when enough cells of that type are available). These gene expression profiles were then summed. To mimic technical variation, Gaussian noise (mean = 0; SD = 0.05) was added to each pseudobulk, and negative expression values were clipped to zero. A total of 500 pseudobulks were generated under each proportion type (realistic and random), yielding 1,000 pseudobulk samples overall. These pseudobulks, for which we had corresponding cell-type ground truth proportion matrices, were used as input to the deconvolution method.

### Held-out cell reference and transformations

For each cell type in our datasets (ADP, PBMC, MBC and MSB), we replaced the scRNA-seq expression with either a “raw” snRNA-seq counterpart, or that same snRNA-seq expression with different transformations (outlined below). We deconvolved the same 1000 pseudobulks with each of the reference types. We also included the positive control (i.e., held-out cell is not replaced, just scRNA-seq).

### Transformations

- **Non-Transformed: Controls (scRNA All (PosCtrl), snRNA All (NegCtrl), snRNA-seq):** In the pseudobulk simulation experiments, we created the following controls to compare the performance of the transformations below. For each dataset, we created one reference that includes all cell types available in the pseudobulk, where all cells come from a scRNA-seq dataset (labeled scRNA All (PosCtrl)). This reference represents the ideal research scenario and positive control, although not realistic in some circumstances. We also created one reference of equal cell types and number of cells per cell type, but where all cells come from a snRNA-seq dataset, labeled (snRNA All (NegCtrl). For the experiments in which we hold-out one scRNA-seq cell, we also created one reference per cell type where remove one cell type at a time from the scRNA All (PosCtrl) reference and replace that cell’s expression with the equivalent from the snRNA All (NegCtrl) reference (labeled snRNA-seq). In the real ADP data experiments with real bulks, using a reference will all cell types in scRNA-seq or all cell types in snRNA-seq without integration would not be comparable; the datasets do not have matched cell types (i.e., one and two cell types missing) even when we integrate multiple datasets from each. We created the “snRNA-seq” reference by combining wall the cells available in scRNA-seq datasets and added the remaining cell types (missing from scRNA-seq) from snRNA-seq datasets, mimicking a real-world scenario.
- **Differentially expressed genes removed (snRNA All (-DEG Int.), snRNA -DEG, - Random Genes, -DEG Other Datasets, -DEG Int.):** To calculate the differentially expressed genes between scRNA-seq and snRNA-seq per cell type, we created S-cell aggregated and snRNA-seq aggregated (sum of the expression of 10 cells) for each cell type as recommended in [34]. We used this cell type aggregates to compute the DEG using pyDESEQ2 version 0.5.0 [35]. DEGs were defined as those with a p-adjusted value of less than 0.01 after Benjamini-Hochberg adjustment. We removed the union of DEGs of all cell types (per dataset) from some of the cell references (labeled snRNA -DEG), therefore removing them from the deconvolution process. We tested the removal of other gene groups in deconvolution. These reference test whether pruning improves alignment of the bulk anf reference, and therefore improve deconvolution. For the snRNA All (-DEG Int.) reference, created only for human datasets, we used all snRNA-seq cells (no held out), and removed the list of genes that was found to be differentially expressed in all human datasets (the intersection or Int.). We additionally created references with a random set of genes removed of the same number as the DEGs (labeled -Random) to validate the biological context of the calculated -DEGs. For the human datasets, we also created references with the intersection DEGs removed (-DEG Int.) (similar to snRNA All (-DEG Int.), but all scRNA-se cells with added snRNA-seq), and removing the DEGs that were found in the other datasets not including the current data (e.g., removing the DEGs from ADP and PBMC in MBC) (labeled -DEG Other Datasets). The details on the references, genes removed, and cells included are outlined in Table 1.
- **PCA neighbour-based shift (PCA and PCA-DEG):** For each dataset (ADP, PBMC, MSB, and MBC), cell counts for scRNA-seq and overlapping cell types in snRNA-seq were first transformed with log (x + 1) and then standardized to zero mean and unit variance. Principal-component analysis was fitted to the scaled data, retaining the minimum number of components required to capture at least 75 % of the total variance. The scRNA-seq and overlapping snRNA-seq cells were projected into this lower dimensional PCA space. We shift the cell’s expression similar to what is done in [36]: we first found the centroid of all scRNA-seq observations. For each (only overlapping) snRNA-seq cell, we calculated an observation-specific shift as the distance between that cell and the scRNA-seq centroid, giving us a list of distances for each overlapping snRNA-seq cell. Then, the held-out or “missing” cell type (for which we do not have a scRNA-seq example) cells were projected into the same fitted PCA space. For each cell example of this “missing” cell type, we calculated the Euclidian distance to other snRNA-seq cells and found the 10 nearest neighbours in the PCA space. We then used the mean distance of these 10 nearest neighbours as a shift vector for each cell, and this shift was added to each cell according to its nearest neighbours, shifting the snRNA-seq expression. Finally, the shifted latent representations were back-projected to gene expression space by applying the inverse PCA transform, followed by taking the natural exponential and then subtracting 1 to revert the log-transformation, and subsequently scaled to match the median library size of the scRNA-seq data (mean count value). Each of these steps was repeated for the datasets with the DEGs removed (labeled PCA -DEG).
- **scVI (VAE) model with latent space shift (scVILS and scVILS -DEG):** For each dataset (ADP, PBMC, MSB, and MBC), we used a traditional scVI VAE model to “transform” one data type (snRNA-seq) to another (scRNA-seq) through alignment in the latent space. We trained multiple scVI models for each data type: one for each removed cell, ensuring the training data never contained the held-out or “missing” cell type to simulate a real life scenario [13]. For all models, we used 2 layers, 30 latent variables, gene-batch dispersion and negative binomial gene likelihood (default parameters otherwise) because these parameters have shown to work well in alignment tasks [25], with no conditional encoding of data type. All models were trained with an early stopping patience of 10 epochs. All code showing model training can be found in the GitHub repository under scripts/ train_scvi_models_allgenes.py and scripts/train_scvi_models_nodeg.py without DEGs [30]. After training, we encoded the snRNA-seq held-out or “missing” cells to the latent space and shifted each cell the same way it was done in the PCA neighbour-based shift described above. We then decoded the shifted expression and used the median library size from the scRNA-seq cells to scale the decoded snRNA-seq data, as is described above for the PCA shift. We repeated this reference with and without the the DEGs (i.e., new models trained with less gene features) as computed above.
- **scVI (VAE) conditional model (scVIcond and scVIcond -DEG):** For each dataset (ADP, PBMC, MSB, and MBC), we used a conditional scVI VAE model to “transform” one data type (snRNA-seq) to another (scRNA-seq). We trained multiple scVI models for each data type: one for each removed cell, ensuring the training data never contained the held-out or “missing” cell type to simulate a real life scenario [13]. All models were set to be conditional on the data type, either scRNA-seq or snRNA-seq, by one-hot encoding the data type label as a feature (encoded at encoder and injected at latent space). For all datasets and cell types, we used 2 layers, 30 latent variables, gene-batch dispersion and negative binomial gene likelihood (default parameters otherwise) because these parameters have shown to work well in alignment tasks [25]. The training parameters were kept constant and the same as described above for scVI (VAE) latent space neighbour-based shift. All code showing model training can be found in the GitHub repository under scripts/ train_scvi_models_allgenes.py and scripts/train_scvi_models_nodeg.py without DEGs [30]. After training, we encoded and decoded the snRNA-seq removed cell with scRNA-seq labels, which we hypothesize would cause the model to transform the snRNA-seq expression into a scRNA-seq counterpart. We used the median library size from the scRNA-seq cells to scale the decoded snRNA-seq data. We repeated this reference with and without the DEGs (i.e., new models trained with less gene features) as computed above.

### Comparison of snRNA-seq transformed cell types with real scRNA-seq cells

For each cell type in each dataset (Figure 1b) we compared the expression profile obtained after every snRNA transformation with the corresponding profile from the scRNA-seq reference. For both modalities, we randomly sampled 50 cells per cell type and aggregated them to reduce sparsity by summing counts to form a pseudobulk vector; the minimum number of cells across all cell types and datasets. We normalized the expression to counts per million and log-transformed to makes samples comparable across library sizes, and log makes the similarity reflect relative expression patterns rather than being dominated by highly expressed genes. By using raw counts, the cosine similarity would be dominated by total read depth, not true biological differences. We then calculated cosine similarity between each transformed snRNA-seq vector and its scRNA-seq counterpart. The procedure was repeated for all transformations, with untransformed snRNA-seq serving as a negative control and scRNA-seq as the positive control. The similarity scores were then summarized across cell types.

### Comparison of transformed cell types with real bulks

We created pseudobulks as outlined in the previous Pseudobulks section (same number of cells and sampling logic). We created 100 realistic-proportioned pseudobulks and 100 random-proportioned pseudobulks for each reference type (PCA, scVILS, scVIcond, and repeated without -DEGs). For all pseudobulks, we used all scRNA-seq cells and added the snRNA-seq “missing” cells (adipocytes and neutrophils) either raw or with each of our transformations listed above. We aimed to see which transformation made the pseudobulks containing the transformed cells be closer to the real bulks by computing the cosine similarity. We first library-normalized the data to counts per million to remove the sequencing depth (or cell number) variation, and log + 1 the data to stabilize the variance. We then computed the cosine similarities. For each transform, we obtained a distribution of cosine similarity scores: one value for every pseudobulk x realbulk comparison. We then computed the 95% bootstrapped CI of the mean with 1000 iterations.

### Deconvolution results and comparison

For deconvolution, we used BayesPrism [10] metholdogy through InstaPrism [37], a probabilistic framework that leverages the expression profiles of the cell reference to deconvolve bulk mixtures that has been shown to outperform others in previous work [27]. We executed the InstaPrism deconvolution for 5000 iterations for each bulk, per reference dataset, including the controls using only the scRNA-seq data (“scRNA All (PosCtrl)”) and only the snRNA-seq data (“snRNA All (NegCtrl)”), as well as the transformed hybrids. The deconvolution output consisted of estimated cell type proportions for each bulk/pseudobulk sample.

In the case of the simulation experiments (i.e., pseudobulks) we have ground truth proportions which we can use to compute performance metrics. We compared the estimated cell-type proportions with the known ground-truth proportions with two quantitative metrics: Pearson correlation to assess the linear concordance between predicted and true proportions, and RMSE to quantify the overall estimation error.

We evaluated performance under three scenarios for each bulk:

- **All Cells:** The evaluation was performed using the entire set of cell types present in the simulated bulk (all proportions of all cells estimated).
- **Non-Removed Cells:** The analysis was repeated after excluding the held-out cell type from the evaluation. This scenario represents the case where the reference already contains the cell types that are present in the bulk.
- **Removed Cell Only:** In a realistic missing-cell scenario, one cell type was intentionally removed from the scRNA-seq reference and replaced with the transformed snRNA-seq data. Performance for this held-out cell type was evaluated separately. For this scenario, controls (i.e., references that did not have any held out any cell type) were excluded since there is no direct comparison (no held-out cell type).

For the simulations, we show the mean performance per transforms along with the 95% bootsrapped CI of the mean with 1000 iterations. For the non-simulation experiments (real bulks deconvolved with each transform type, and with each patient’s datasets), we do not have ground truth proportions, so we evaluate the robustness as outlined below (see Real adipose bulks deconvolution robustness).

For the comparison of using scRNA-seq vs. snRNA-seq as reference in pseudobulks made from scRNA-seq cells, we also conducted independent two sample Student’s t test to compare the mean performance metrics between scRNA-seq reference and snRNA-seq reference across each of the datasets. We tested for differences in the Pearson correlation coefficients and the RMSE deconvolution performance values by performing two tailed t-tests (assuming equal variances), with any missing values omitted on a pairwise basis. Statistical significance was evaluated at the 0.005 level.

### Gene Ontology analysis of intersection genes

We identified a list of genes that is differentially expressed in at least one cell type across all 3 human datasets (ADP, PBMC, MBC). We used GOrilla (Gene Ontology enRIchment anaLysis and visuaLizAtion tool) [38] to test for over-representation of GO Cellular Component terms in the intersection gene set. The “target” list comprised the “intersection genes”, and the “background” list comprised every gene detectable in all three of the datasets (ADP, PBMC, MBC). Enrichment was calculated with the two-list (target vs. background) hypergeometric test implemented in GOrilla, and p-values < 0.05 were considered significant. We considered the first 30 components for visualization in Figure S3.

### Real adipose bulks data and single modality data preprocessing

Bulk RNA seq data from 331 subcutaneous adipose tissue samples (METSIM cohort, GEO GSE135134 [20]) were processed by first importing TPM expression values from the publicly available downloaded file (see Data Availability). Gene-level annotation was performed using a Gencode v47 basic GTF file [39]. Finally, only the genes common to the bulk data and corresponding snRNA-seq and scRNA-seq datasets were retained, and the resulting expression matrix was saved for downstream analyses.

For the accompanying scRNA-seq and snRNA-seq datasets, we used the same data source [20] (not same datasets) as used for the adipose data in the deconvolution experiment. For this experiment, we filtered to only datasets that are the same tissue type as the real bulks (subcutaneous adipose tissue) that were not used as references or for deconvolution in the previous experiment. This yielded 7 scRNA-seq and 12 snRNA-seq patient datasets. We filtered each of the datasets independently. We added a table with all data links and processing parameters for each dataset to the GitHub repository (/data/details/Data_Details.xlsx) [30].

### Real Bulks Deconvolution Robustness (per transform and per patient)

We used deconvolution, as described above, to estimate the proportions of cell types in 434 real bulk samples. We used the adipose tissue datasets as references (7 scRNA-seq datasets, 11 snRNA-seq datasets). These datasets had two additional cell types in snRNA-seq that were not present in any scRNA-seq dataset (fat cells and neutrophils), and one cell in scRNA-seq not present in snRNA-seq, making them an ideal dataset to test what happens in a true research setting. We created a reference with all scRNA-seq cells, and the two additional cells from snRNA-seq, either raw as a control or transformed with each of our transforms (described above).

Previous work has described robustness (i.e., consistency in predictions) as a measure of performance [27] in deconvolution. We did two experiments to evaluate the performance of our transformations; we compared the predicted proportions of each transformation to the predicted proportions of the other transformations. We also tested the robustness of our transformations by comparing the predicted proportions if we only use fat cells from one patient at a time (i.e., inter patient robustness). We created references with all scRNA-seq cell types from all patients, snRNA-seq neutrophils from all patients (these cells have low quantities in all patients, so we pooled them), and snRNA-seq fat cells from each patient at a time transformed with each of our transforms.

We computed the cosine similarity between each transformation’s predicted proportions as two long vectors (transform to transform robustness), and we computed the cosine similarity between each patient’s predicted proportions per transform (e.g., patient 1 cells PCA transform vs. patient 2 cells PCA transform). Finally, we compare the cosine similarities for each transform, both across transformations and across patients. We plot each in x and y axes respectively (Figure 3b).

### Composite score for each transform

We used the RMSE and Pearson correlation values from the simulation experiments (described above), along with the robustness/consistency measures per patient and per transform (described above) to get a new composite score per transform.

For each simulation dataset (ADP, PBMC, MSB, MBC), we took every held-out-cell evaluation (per-sample Pearson correlation and RMSE) and normalized each metric to the range [0,1] using min-max scaling, applied separately within each dataset. For RMSE, we first inverted the values by subtracting them from the dataset maximum (equivalent to multiplying by -1 before scaling), so that higher values indicate better accuracy. We then averaged the normalized Pearson and RMSE values across all samples belonging to the same transform within that dataset. Next, we collapsed across datasets by taking the mean of each transform’s per-dataset normalized Pearson and normalized RMSE. Finally, we defined a single “accuracy” score for each transform as the arithmetic mean of its two normalized metrics. This yields one score that summarizes the RMSE and Pearson correlation across all held-out cell types.

For each reference transformation, we quantified its consistency along two orthogonal axes (across donors and across transforms) and then combined them into a single “consistency score.” First, from the pair-wise donor-to-donor similarity table (all cosine similarities between every two donors’ predicted proportion vectors for a given transform), we min-max scaled those cosine values to [0, 1] and averaged them to yield a per-donor score for each transform. Second, from a transform-to-transform cosine similarity matrix (excluding self comparisons), we likewise flattened the off-diagonal entries into a long list, applied the same min-max scaling, and averaged per transform to obtain a per-transform score. Finally, we defined the overall “robustness score” as the simple mean of the per-donor score and per-transform score, thereby giving equal weight to between-donor and between-reference agreement. This yields one score that summarizes the cosine similarities across transforms and donors.

This gave us two normalized values per transform; one that shows the accuracy seen in pseudobulk experiment (Figure 3e y axis), and one that shows the robustness in real data (Figure 3e x axis).

## Data Availability

The data that support the findings of this study are all freely available online. The count data for ADP and MBC can be downloaded through the Gene Expression Omnibus with accession numbers GSE176067 (adipose single-cell) [40], GSE176171 (adipose single-nucleus) [41], adipose bulks (GSE174475) [42], and GSE140819 (metastatic breast cancer) [43]. The PBMC and MSB data provided by 10x Genomics can be directly accessed through their website [44]. All data details and download links can be easily accessed in the GitHub repository [30].

## Code Availability

The code developed for this study is available at our GitHub repository [30] under BSD 3-Clause License, and a Zenodo repository will be created once article is accepted. The repository includes all necessary scripts and a README file with setup instructions and usage guidelines. For updates or assistance, users can refer to the repository or open an issue for queries. Our aim is to support transparency and reproducibility in computational research through this open-access resource.

