## Supplementary Figures for "Integrating single-cell and single-nucleus datasets improves bulk RNA-seq deconvolution"

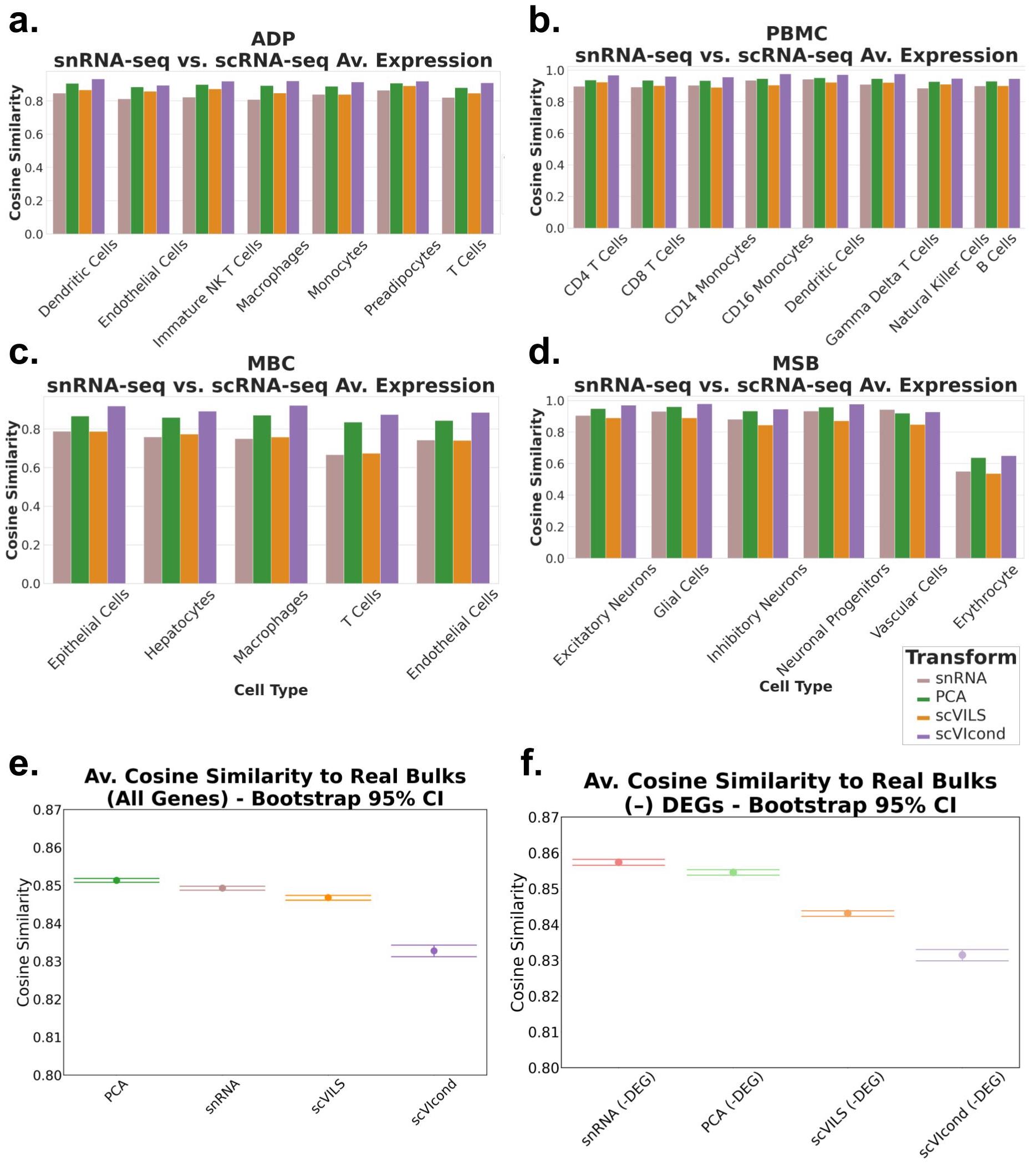

**\**

**Figure S1**

**scVIcond Maximizes Cell-Level Alignment, Whereas PCA-Transformed snRNA-Seq Best Recapitulates Bulk Expression**

Comparison of transformed snRNA-seq data with scRNA-seq and with real bulks. a-d, The cosine similarity between the average expression of ScRNA-seq cells, per cell type, and the average expression of the transformed S-Nuc cells of that same cell type (a: ADP, b: PBMC, c: MBC, d: MSB datasets). e-f, Pseudobulks created from adipose tissue data with all ScRNA-seq cells, and the 2 cells not present in ScRNA-seq were added from snRNA-seq data. The snRNA-seq cells were either control (raw snRNA) or transformed, or all cells from snRNA-seq as a control. e and f. Plots show the mean cosine similarity (dots) of each of these pseudobulks with each of 434 real bulks, expected to have the same cell types. The bootstrapped 95% confidence interval are shown as bars. Y‑axes are truncated to highlight the variation.

ScRNA-seq: single-cell RNA-seq. S-Nuc: single-nucleus RNA-seq, ADP: adipose tissue data. PBMC: peripheral blood mononuclear cells. MBC: metastatic breast cancer. MSB: mouse E18 brain.

**Figure S1 Results Details**

**Cell-type expression alignment shows scVI conditional model best transforms snRNA-seq toward scRNA-seq profiles**

We then aimed to find ways in which we can incorporate cells from snRNA-seq datasets that might not be present in scRNA-seq datasets without compromising performance. We explored different transformations or ways of integrating the snRNA-seq cells into a scRNA-seq reference (described in the Methods section). We first aimed to explore whether each transformed snRNA-seq cell type would resemble a scRNA-seq counterpart more when compared to the original snRNA-seq expression, per cell type, across four datasets (Fig. 1b).

We transformed each cell type in each of the datasets (Fig. 1b) with the scVILS, scVIcond and PCA neighbor-based transforms. We computed the mean expression for each transform in each cell type, along with the original “raw” snRNA-seq expression and the same cell type from scRNA-seq dataset. We calculated the cosine similarity between the target scRNA-seq mean expression and each of our transforms and control (snRNA-seq unchanged).

All gene expression was CPM‑normalized and log‑transformed, so scale and library size differences across protocols are largely removed; what remains are relative gene‑expression patterns. Cosine similarity directly compares these patterns by measuring the angular alignment of two expression vectors, providing a unit‑free score (1 = identical, 0 = orthogonal) that captures how well the transformed S‑Nuc profile follows the shape of the S‑Cell signature, independent of residual shifts in overall amplitude. This makes it an interpretable metric for evaluating transformation quality.

In nearly every cell type across the 4 datasets, the cosine similarity, between scRNA-seq and snRNA-seq was the lowest, suggesting the transformations are shifting the snRNA-seq expression towards a scRNA-seq equivalent (Figure S1a). Overall, the cosine similarity between the scRNA-seq and the scVIcond expression was highest, with PCA as a close second. The cosine similarity for the scVI LS transformation was comparable to the snRNA-seq control, and for some cell types it was lower than the snRNA-seq control, particularly in the MSB data. We suspect this drop reflects a mismatch between a linear neighbor-based adjustment and the nonlinear geometry of the scVI latent space.

**PCA-transformed snRNA-seq pseudobulks best approximate bulk RNA-seq profiles**

We then explored whether the transformed snRNA-seq data would also resemble the expression we would expect to see in real bulk data. We hypothesize that bulk RNA-seq datasets also contain all nuclear and RNA as seen in a scRNA-seq dataset, and that the transformation of snRNA-seq cells would thus improve the deconvolution of real bulks. To be able to compare our transforms (in single-modality scale) to real bulk (aggregated expression), we recreated bulk expression by making pseudobulks using the snRNA-seq transformed cells along with non-transformed cells we would expect to be present in bulk.

We used all cell types available in seven adipose tissue scRNA-seq datasets – which we hypothesize contain all RNA (cytoplasmic and nuclear). The scRNA-seq datasets were missing two cell types when compared to twelve adipose tissue snRNA-seq datasets (fat cells and neutrophils) from the same study, we added the two missing cells’ expression- which we hypothesize are present in real bulk data – transformed with each of the transformations. We created 100 pseudobulks of random proportions and 100 pseudobulks of realistic proportions (as described in Methods: Pseudobulks) for each transformation (i.e., fat cells and neutrophils are transformed, scRNA-seq cells aggregated as is). We also included a control in which the missing cell types are not transformed (snRNA-seq raw expression added to scRNA-seq). We kept the expression (cell examples) constant across all pseudobulks. This design isolates the effect of each transformation while providing an appropriate comparison to bulk RNA-seq data expression.

In real gene space, we normalized the data and calculated the cosine similarity between each pseudobulk and each real bulk (see Methods). We calculated the mean cosine similarity per pseudobulk for all transformations and show the mean and the bootstrapped 95% confidence interval of the mean in Figure S1e. We found only the PCA transformation pseudobulks had a higher cosine similarity to the real bulks when compared to snRNA-seq raw expression pseudobulks. Surprisingly, scVIcond pseudobulks had the lowest cosine similarity.

We repeated this independently for transforms that include all genes and transforms with gene removal to keep a comparable feature space. The transformations labeled -DEG have the genes that were found to be differentially expressed between each cell type in scRNA-seq with that same cell type in snRNA-seq. For details on DEG calculations, see Methods. In this analysis we included all DEGs found between cell types available in both scRNA-seq and snRNA-seq (not fat cells or neutrophils). In the transformations that involve gene filtering, the snRNA-seq data with the removed DEGs had the highest cosine similarity, followed by the PCA -DEG transform (Figure S1f).

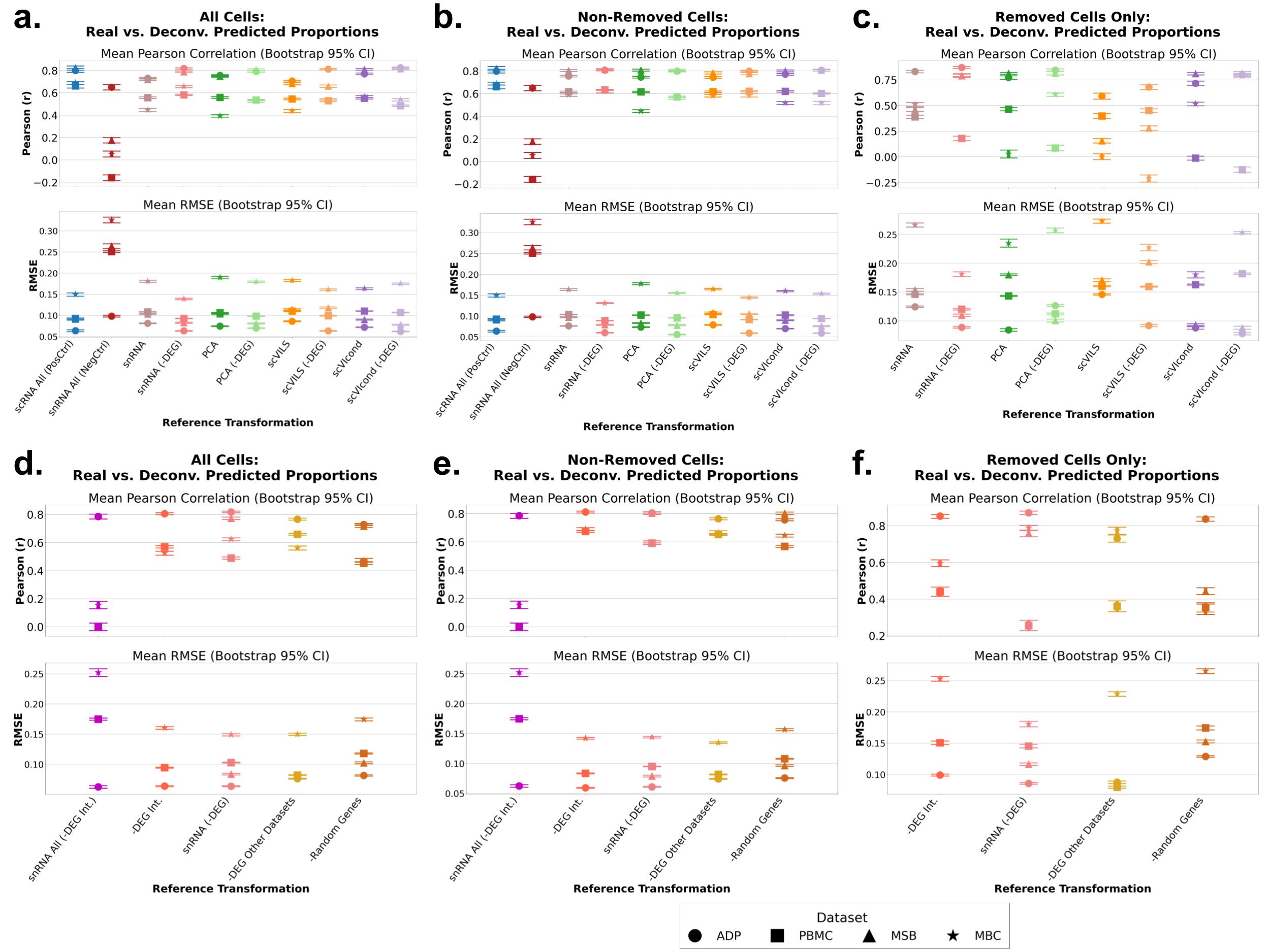

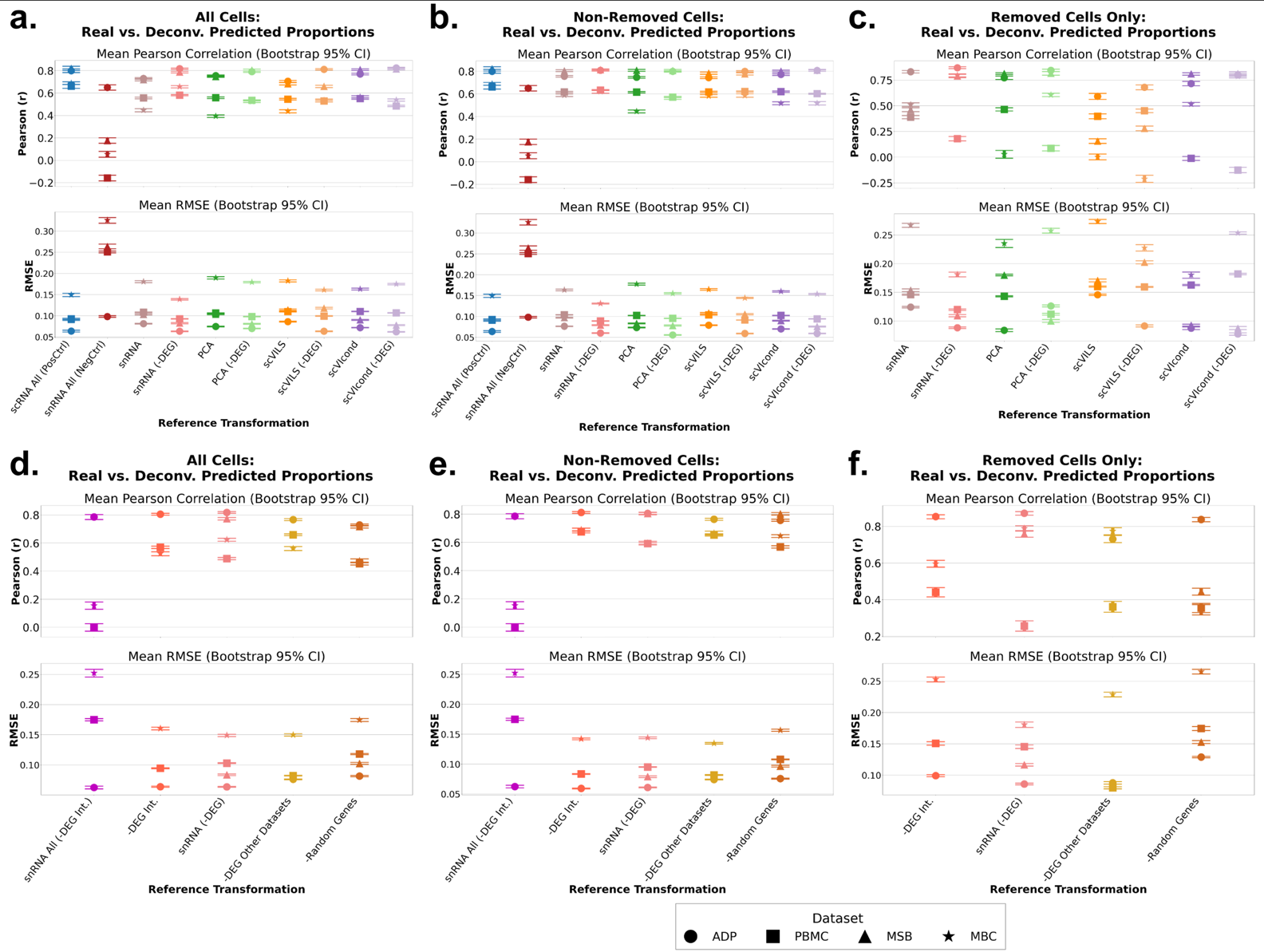

**Figure S2**

**Pseudobulk deconvolution accuracy with each cell type in scRNA-seq held out and transformed.**

Each plot shows the Pearson correlation value (top panels) and the RMSE values (bottom panel) for the ground truth pseudobulk (simulated) proportions and the predicted proportions. We hold out one cell type at a time from each dataset and replace that cell type’s expression with a snRNA-seq equivalent with each of the transformations or controls (scRNA-seq All and snRNA-seq All) on the x axis of each plot. Each dot represents the mean metric (r or RMSE) across datasets (Figure 1b) and the bars represent the 95% bootstrapped confidence interval of the mean. We evaluated 3 scenarios (see Methods for details). a and d. All cells included in performance metrics calculations, b and e. Non-Removed cells only included in performance metrics calculations and c and f. Only removed cells included in performance metrics calculations. Y‑axes are truncated to highlight the variation. Combined metrics for all datasets are shown in Figure 2.

scRNA-seq: single-cell RNA-seq, RMSE: root mean squared error, snRNA-seq: single-nucleus RNA-seq.

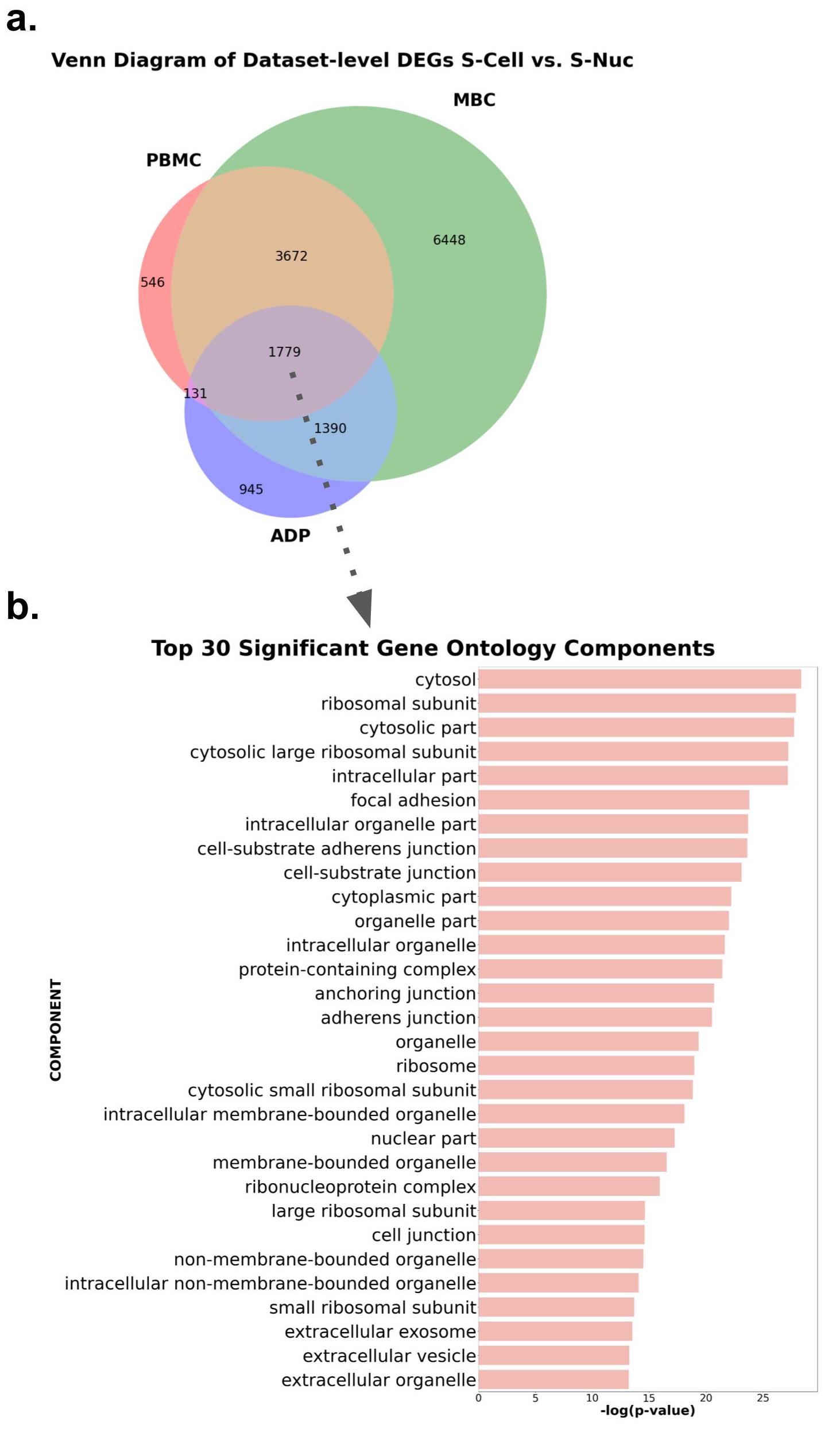

**Figure S3**

**Shared and unique scRNA-seq vs snRNA-seq differentially expressed genes across three human datasets, with GO-component enrichment of the common signature.**

DEGs between scRNA-seq and snRNA-seq cells of the same cell types in the three human datasets. a. Venn diagram showing the number of DEGs in common and distinct between cell types. b. Significant (GOrilla default of p <0.005) gene ontology terms (component) identified from the intersection genes (target) compared to the full gene list (all genes in common between datasets).

DEG: differentially expressed gene. ScRNA-seq: single-cell RNA-seq. snRNA-sequencing: single-nucleus RNA-sequencing. ADP: adipose tissue. PBMC: peripheral blood mononuclear cells. MBC: metastatic breast cancer.

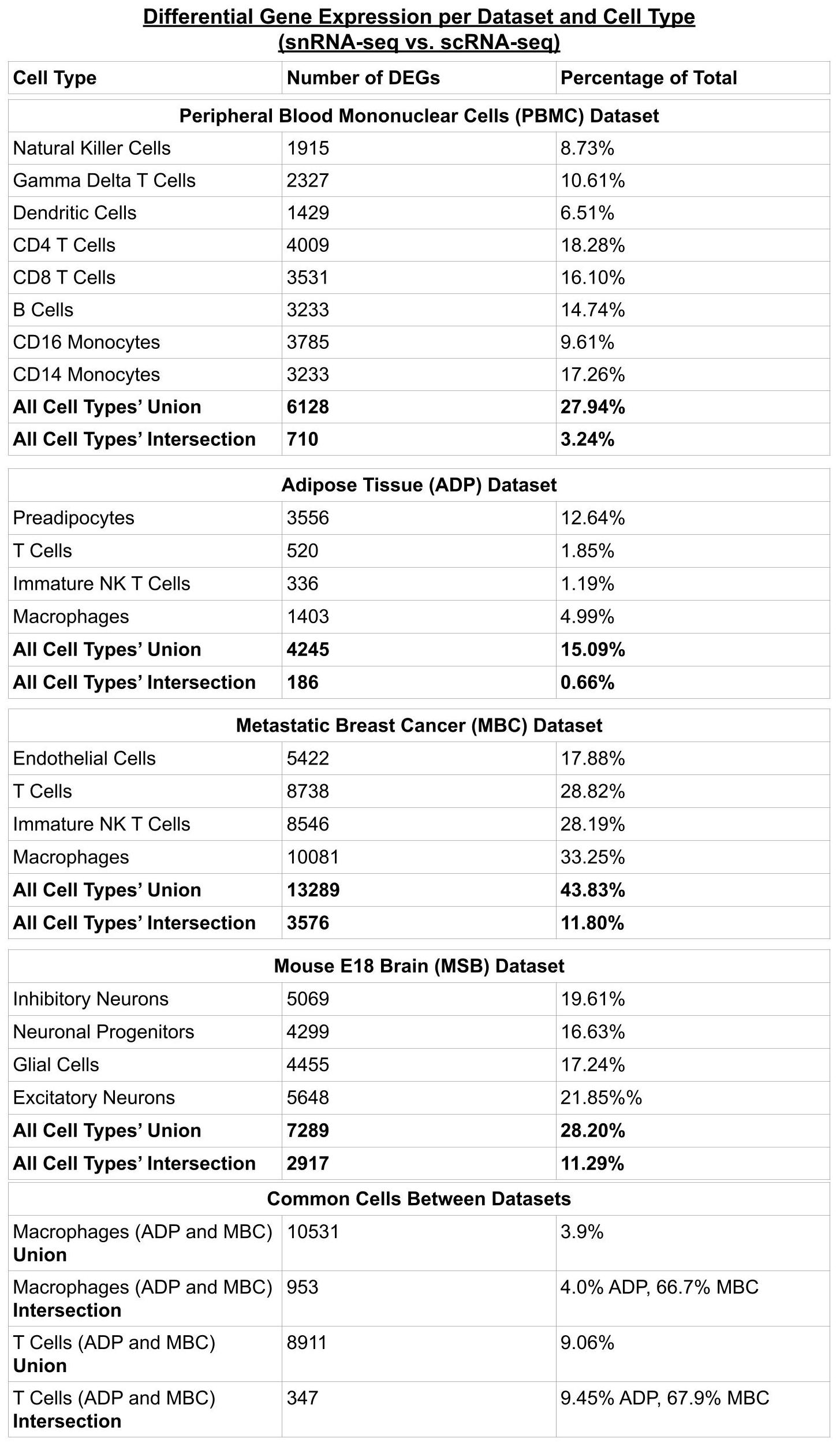

**Table 2.**

**Supp. Table 1**

**Number of union and intersection genes per cell type in each of the datasets**. First row (left) shows the cell type, middle row shows the number of DEGs between that cell type in scRNA-seq vs. S-Nuc. The right-most row shoes the percentage of total genes that are classified as DEGs. For each dataset, we note the number of genes that are in common between the cell types, both union (all) and intersection (common between all cell types). The human datasets contained 2 cell types in common across datasets, and we note the union and intersection of the DEGs between datasets as well.

DEG: differentially expressed gene. scRNA-seq: single-cell RNA-seq. snRNA-seq: single-nucleus RNA-seq.
